# Codon usage and evolutionary rates of the 2019-nCoV genes

**DOI:** 10.1101/2020.03.25.006569

**Authors:** Maddalena Dilucca, Sergio Forcelloni, Athanasia Pavlopoulou, Alexandros G. Georgakilas, Andrea Giansanti

## Abstract

Severe acute respiratory syndrome coronavirus 2 (2019-nCoV), which first broke out in Wuhan (China) in December of 2019, causes a severe acute respiratory illness with a mortality ranging from 3% to 6%. To better understand the evolution of the newly emerging 2019-nCoV, in this paper, we analyze the codon usage pattern of 2019-nCoV. For this purpose, we compare the codon usage of 2019-nCoV with that of other 30 viruses belonging to the subfamily of orthocoronavirinae. We found that 2019-nCoV shows a rich composition of AT nucleotides that strongly influences its codon usage, which appears to be not optimized to human host. Then, we study the evolutionary pressures influencing the codon usage and evolutionary rates of the sequences of five conserved genes that encode the corresponding proteins (viral replicase, spike, envelope, membrane and nucleocapsid) characteristic of coronaviruses. We found different patterns of both mutational bias and nature selection that affect the codon usage of these genes at different extents. Moreover, we show that the two integral membrane proteins proteins (matrix and envelope) tend to evolve slowly by accumulating nucleotide mutations on their genes. Conversely, genes encoding nucleocapsid (N), viral replicase and spike proteins are important targets for the development of vaccines and antiviral drugs, tend to evolve faster as compared to other ones. Taken together, our results suggest that the higher evolutionary rate observed for these two genes could represent a major barrier in the development of antiviral therapeutics 2019-nCoV.

## 1. Introduction

The name “coronavirus” is derived from the Greek *κ*o*ρωνα*, that mean crown. The first complete genome of coronavirus (mouse hepatitis virus - MHV), a positive sense, single-stranded RNA virus, was first reported in 1949. It belongs to the family Coronaviridae and ranges from 26.4 (ThCoV HKU12) to 31.7 (SW1) kb in length [10], making it the largest genomes among all known RNA viruses, with G + C contents varying from 32% to 43% [20]. The coronavirinae family consists of four genera based on their genetic properties, including genus: *Alphacoronavirus, Betacoronavirus* (subdivided in subgroups A, B, C and D), *Gammacoronavirus* and *Deltacoronavirus*. Coronavirus can infect humans and many different animal species, including swine, cattle, horses, camels, cats, dogs, rodents, birds, bats, rabbits, ferrets, mink, snake, and other wildlife animals. To date, 30 CoVs genomes have been identified. Only seven CoVs have been identified that infect humans, including Human CoV-229E (HCoV-229E), Human coronavirus NL63 (HCoV-NL63), HumanCoV-OC43 (HCoV-OC43), Human CoV-HKU1 (HCoV-HKU1), Human SARS related coronavirus (SARSr-CoV), Middle East Respiratory Syndrome (MERS-CoV) and SARS CoV 2 (nCoV) [26].

Severe acute respiratory syndrome CoV 2 (2019-nCoV), which first broke out in Wuhan (China) in December of 2019, causes a severe acute respiratory illness with a mortality rate from 3% to 6%. The newly sequenced virus genome encodes two open reading frames (ORFs), ORF1a and ORF1ab, the latter encodes replicase polyproteins, and four structural proteins [19], [9]: the spike-surface glycoprotein (protein S), the small envelop protein (protein E), the matrix protein (M), and the nucleocapsid protein (N).

The phenomenon of codon usage bias (CUB) exists in many genomes including RNA genomes and it is actually determined by mutation and selection [1]. As it is well-known, the non-random choice of synonymous codons varies among species that are potential host for viruses [11] [**?**]. It is then important to study patterns of common codon usage in coronaviruses because CUB can be related to the driving forces that shape the evolution of small RNA viruses. Mutational bias has been considered as the major determinant of codon usage variation among RNA viruses [13]. Indeed, RNA viruses show an effective number of codons (ENC) that is quite high (ENC>45) pointing to a quite random codon usage, whereas if one uses the adaptive index CAI then the viral CUB is consistent with that of the host, as observed in the Equine infectious anemia virus (EIAV) or Zaire ebolavirus (ZEBOV) [5]. The aim of this study was to carry out a comprehensive analysis of codon usage and composition of coronavirus and explore the possible leading evolutionary determinants of the biases found.

## 2. Results

### 2.1. Nucleotide composition

The nucleotide content of 30 coronaviruses coding sequences are calculated. The results reveal that the nucleotide A is the most frequent base and the nucleotide frequencies are in order A >T>G>C [19]. For nCoV we note a different trend T>A>G>C, but in totally AT% is more used than GC% (See Table 3). The GC content in 2019-nCoV is 0.373 *±* 0.05.

**Table 1.**
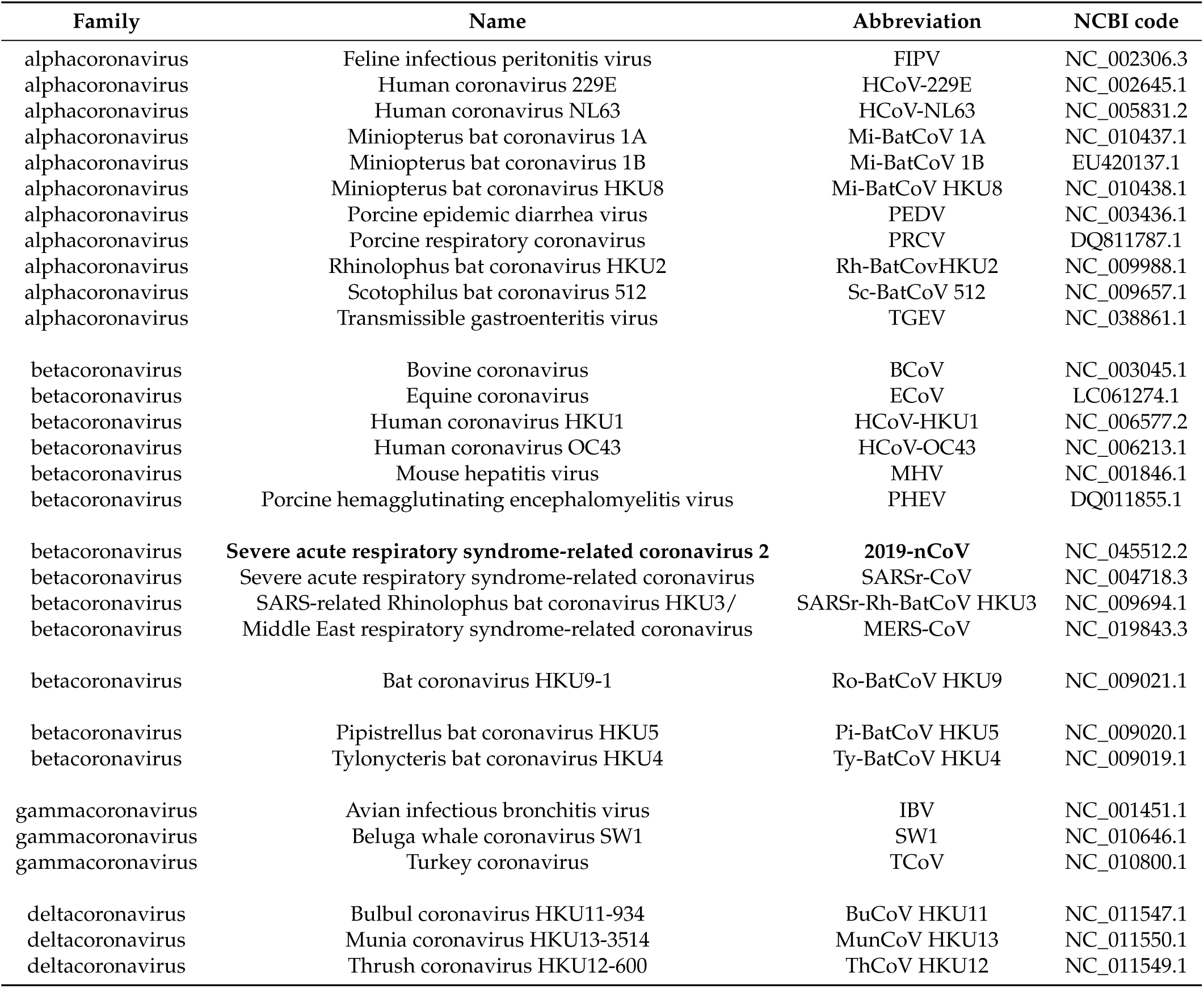
Coronaviruses (CoVs) in dataset. We report name, abbreviation, NCBI code and bp size for each virus.

**Table 2.** Codon usage bias of different coronavirus.

| Abbr. | ENC | CAI |
| --- | --- | --- |
| BCoV | 52.089 ± 2.357 | 0.690 ± 0.040 |
| BuCoV HKU11 | 51.414 ± 1.855 | 0.679 ± 0.045 |
| ECoV | 49.311 ± 4.024 | 0.691 ± 0.025 |
| FIPV | 51.568 ± 1.999 | 0.667 ± 0.048 |
| HCoV-HKU1 | 44.582 ± 7.331 | 0.675 ± 0.021 |
| HCoV-229E | 50.290 ± 3.629 | 0.684 ± 0.023 |
| HCoV-NL63 | 44.675 ± 5.350 | 0.664 ± 0.028 |
| HCoV-OC43 | 49.573 ± 3.660 | 0.692 ± 0.020 |
| IBV | 50.654 ± 2.900 | 0.653 ± 0.053 |
| MERS-CoV | 53.087 ± 2.526 | 0.689 ± 0.029 |
| MHV | 53.628 ± 1.725 | 0.712 ± 0.023 |
| Mi-BatCoV 1A | 48.233 ± 3.810 | 0.682 ± 0.028 |
| Mi-BatCoV 1B | 49.311 ± 4.116 | 0.681 ± 0.026 |
| Mi-BatCoV HKU8 | 50.120 ± 4.139 | 0.702 ± 0.016 |
| MunCoV HKU13 | 53.963 ± 0.863 | 0.691 ± 0.041 |
| **2019-nCoV** | 51.984 ± 2.596 | 0.727 ± 0.054 |
| PEDV | 52.442 ± 2.153 | 0.683 ± 0.037 |
| PHEV | 51.093 ± 3.553 | 0.684 ± 0.024 |
| Pi-BatCoV HKU5 | 53.913 ± 1.359 | 0.691 ± 0.042 |
| PRCV | 51.276 ± 3.154 | 0.675 ± 0.029 |
| Rh-BatCoV HKU2 | 48.084 ± 4.491 | 0.693 ± 0.016 |
| Ro-BatCoV HKU9 | 50.910 ± 2.311 | 0.686 ± 0.033 |
| SARSr-CoV | 53.642 ± 2.436 | 0.670 ± 0.038 |
| SARSr-Rh-BatCoV HKU3 | 54.305 ± 1.613 | 0.667 ± 0.030 |
| Sc-BatCoV 512 | 52.379 ± 2.631 | 0.680 ± 0.043 |
| SW1 | 50.865 ± 1.791 | 0.706 ± 0.027 |
| TCoV | 51.343 ± 2.317 | 0.659 ± 0.046 |
| TGEV | 51.393 ± 3.269 | 0.665 ± 0.037 |
| ThCoV HKU12 | 51.432 ± 2.833 | 0.681 ± 0.031 |
| Ty-BatCoV HKU4 | 50.374 ± 3.742 | 0.683 ± 0.032 |

**Table 3.**
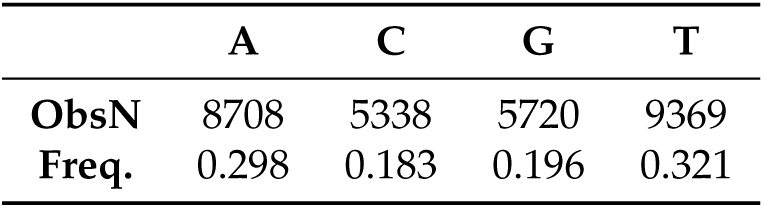
Statistics in nCoV.

|  | A | C | G | T |
| --- | --- | --- | --- | --- |
| **ObsN** | 8708 | 5338 | 5720 | 9369 |
| **Freq.** | 0.298 | 0.183 | 0.196 | 0.321 |

### 2.2. All the sequenced 2019-nCoV genomes share a common codon usage

We downloaded the protein-coding sequences of 2019-nCoV from GIDAID database, by classifying each n-CoV based on the geographic location where they are sequenced (see tree in A2). For each n-CoV genome, we calculated the relative synonymous codon usage (RSCU), in the form of a 61-component vector. In Fig A1, we show the heatmap and the associated clustering of these vectors. Despite the high mutation rate that characterizes 2019-nCoV genome [30], [4], the overall codon usage bias among 2019-nCoV strains appears to be similar. Moreover, their associated vectors did not cluster according to geographic location, thus confirming the common origin of these genomes. Therefore, we will consider a unique vector to represent 2019-nCoV in the following analyses.

### 2.3. Codon usage of 2019-nCoV genome strains

We compared the codon usage of 2019-nCoV with that of the other coronavirus genomes. For this purpose, we used the RSCU which is a biologically relevant metric of distance between the codon usage in the protein-coding sequences of these genomes. In Fig. 1, we report the heatmap of the RSCU values associated with the coronavirus. The RSCU values of the majority of the codons scored between 0 and 3.1 (see legend in Figure 1). Interestingly, analysis of codon bias in coronavirus genomes reveal that the newly identified coronavirus 2019-nCoV Wuhan-Hu-1 sequence has a clear origin from other beta-CoV’s and is closer, specifically, to SARSr-CoV, consistent with their phylogenetic relationships (Figure A3). In line with previous observations, we show that the mean CpG relative abundance in the coronavirus genomes is markedly suppressed [27]. Specifically, GGG, GGC, CCG (pyrimidine-CpG) and ACG (purine-CpG) present a very low frequence of occurrence, probably due to the relative tRNA abundance of the host. In fact, in human the abundance of tRNA in codons GGG, GGC, CCG and ACG are respectavely: 0, 0, 4 and 7.

**Figure 1.**
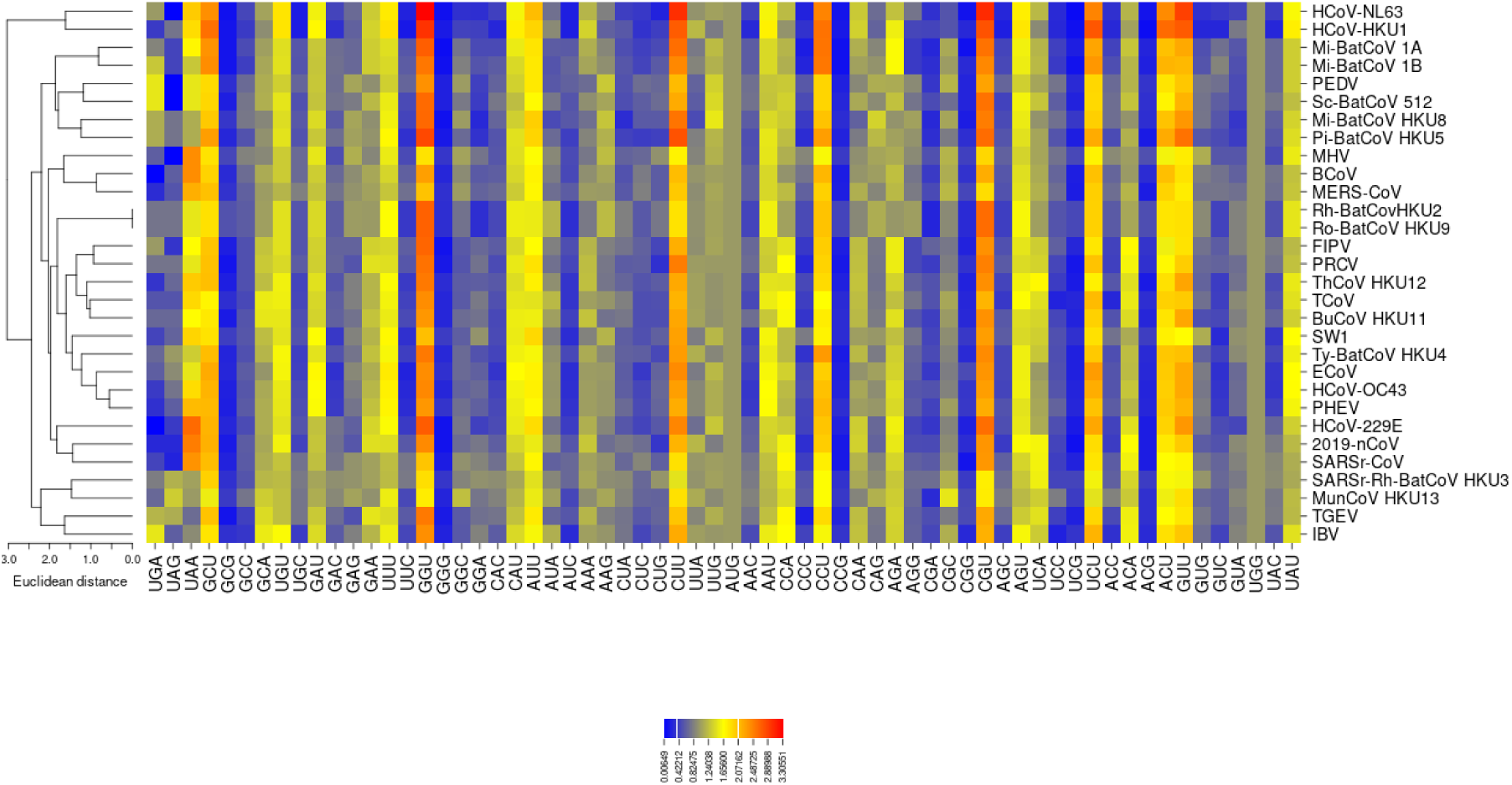
Clustering of the RSCU vectors associated to coronavirus. Analysis of codon bias in coronavirus genomes reveal that the newly identified coronavirus 2019-nCoV Wuhan-Hu-1 sequence was closer to SARSr-CoV as in the reconstructed phylogenetic tree shown in Figure A3. Heatmap are drawn with the CIMminer software [25], which uses Euclidean distances and the average linkage algorithm.

**Figure 2.**
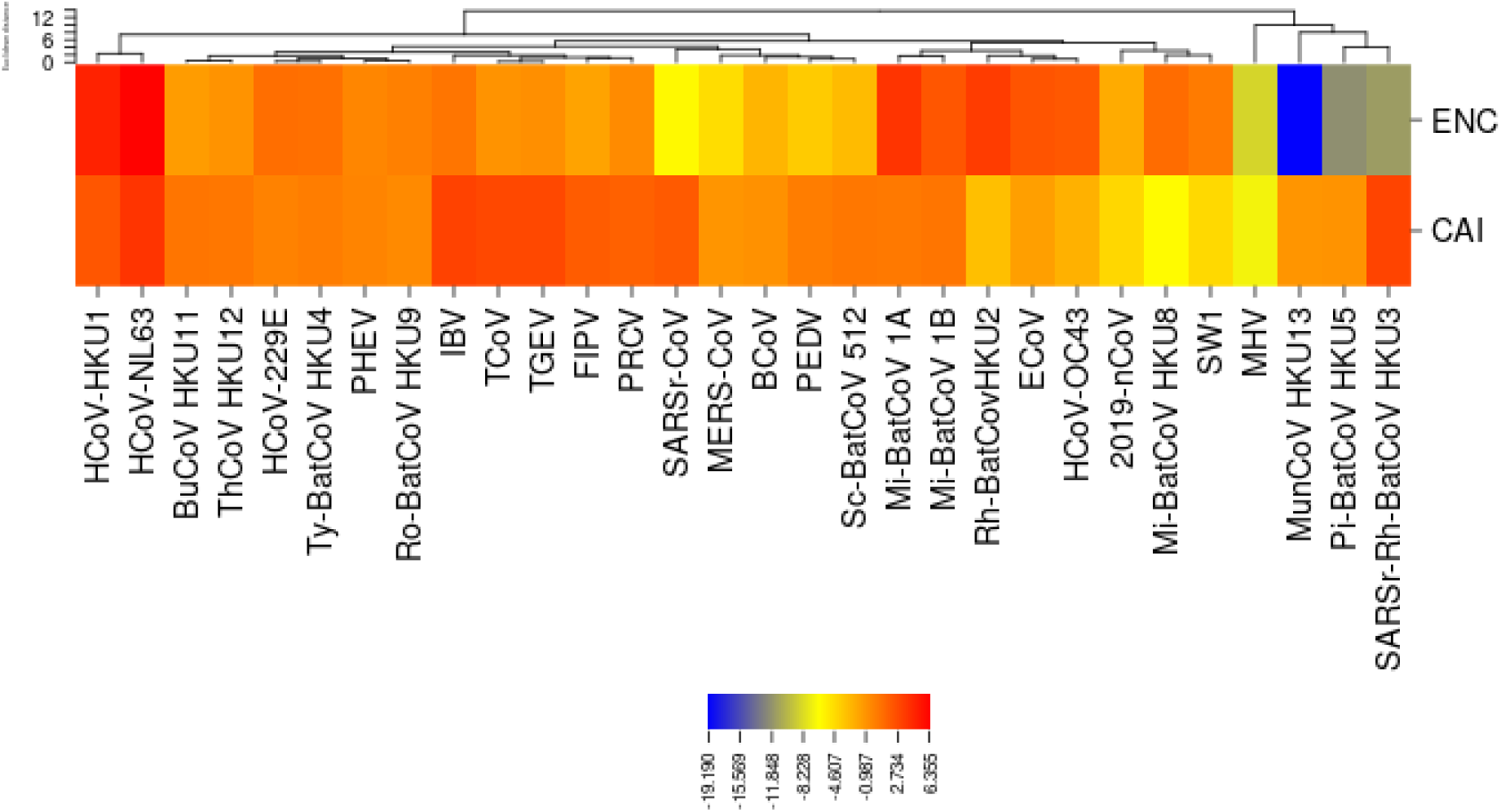
Z-score values. Z-score is calculated for two codon bias indexes ENC and CAI. We show in figure that a lot of coronavirus have a codon usage major than the average value of the family (Z-score 3). For what concerns 2019-nCoV, it presents average CAI and ENC higher than the average too.

In 2019-nCoV, the most frequently used codons are CGU (arginine, 2.34 times) and GGU (glycine, 2.42), whereas the least used codons are GGG (glycine) and UCG (serine). We also note the most frequently used codons for each amino acid, ended in either U or A. [5]

### 2.4. The codon usage of 2019-nCoV is not optimized to the host

To measure the codon usage bias in the coronavirus genomes, we used the effective number of codons (ENC) value and the Competition Adaptation Index (CAI). In Table 2 and 3, we report the ENC and CAI values for all the coronaviruses we considered in this work. To visually enhance the differences among the codon usage of these coronaviruses we calculated Zscore of a single virus in relation to the other members of the CoV subfamily. Interestingly, the ENC value associated to 2019-nCoV (51.9 *±* 2.59) is lower than the average of all Coronaviruses, meaning that 2019-nCoV uses a broader set of synonymous codons in its coding sequences. We also note that CAI of 2019-nCoV (0.727 *±* 0.054) is lower than the average one, underscoring that this coronavirus tend to use codons that are not optimized to the host. We calculate similarity index (SiD) to measure the effect of codon usage bias of the host (human) on the 2019-nCoV genes. The SiD values range from 0 to 1, with higher values indicating that the host has a dominant effect on the usage codons. [16] In our case SiD for 2019-nCoV is equal to 0.78.

### 2.5. Genes

The genome of the newly emerging 2019-nCoV consists of a single, positive-stranded RNA, that is approximately 30K nucleotides long. The overall genome is organizated similar to other coronaviruses. Ceraolo et al. performed a cross-species analysis for all proteins encoded by the 2019-nCoV (see Figure 3-4 in [3]). The newly sequenced virus genome encodes polyproteins, common to all betacoronavirus, which are further cleaved into the individual structural proteins E, M, N and S, as well the non-structural RdRP [28]. So, only five viral genes are classified according to their viral location and studied for each virus because the short length and insufficient codon usage diversity of the other genes might have biased the results. The corresponding gene products including S protein regulates virus attachment to the receptor of the target host cell [2]; the E protein functions to assemble the virions and acts as an ion channel [18], the M protein plays a role in virus assembly and is involved in biosynthesis of new virus particles [17] the N protein forms the ribonucleoprotein complex with the virus RNA [19]. Finally, RdRP catalyzes viral RNA syntesis. We calculate for these five proteins the RSCU vectors in each virus of the dataset (see Figure 3). We show that only for gene E, M and N 2019-nCoV cluster with SARSr-CoV and SARSr-Rh-BatCoV HKU3, consistent with the inferred phylogeny shown in Figure A3.

**Figure 3.**
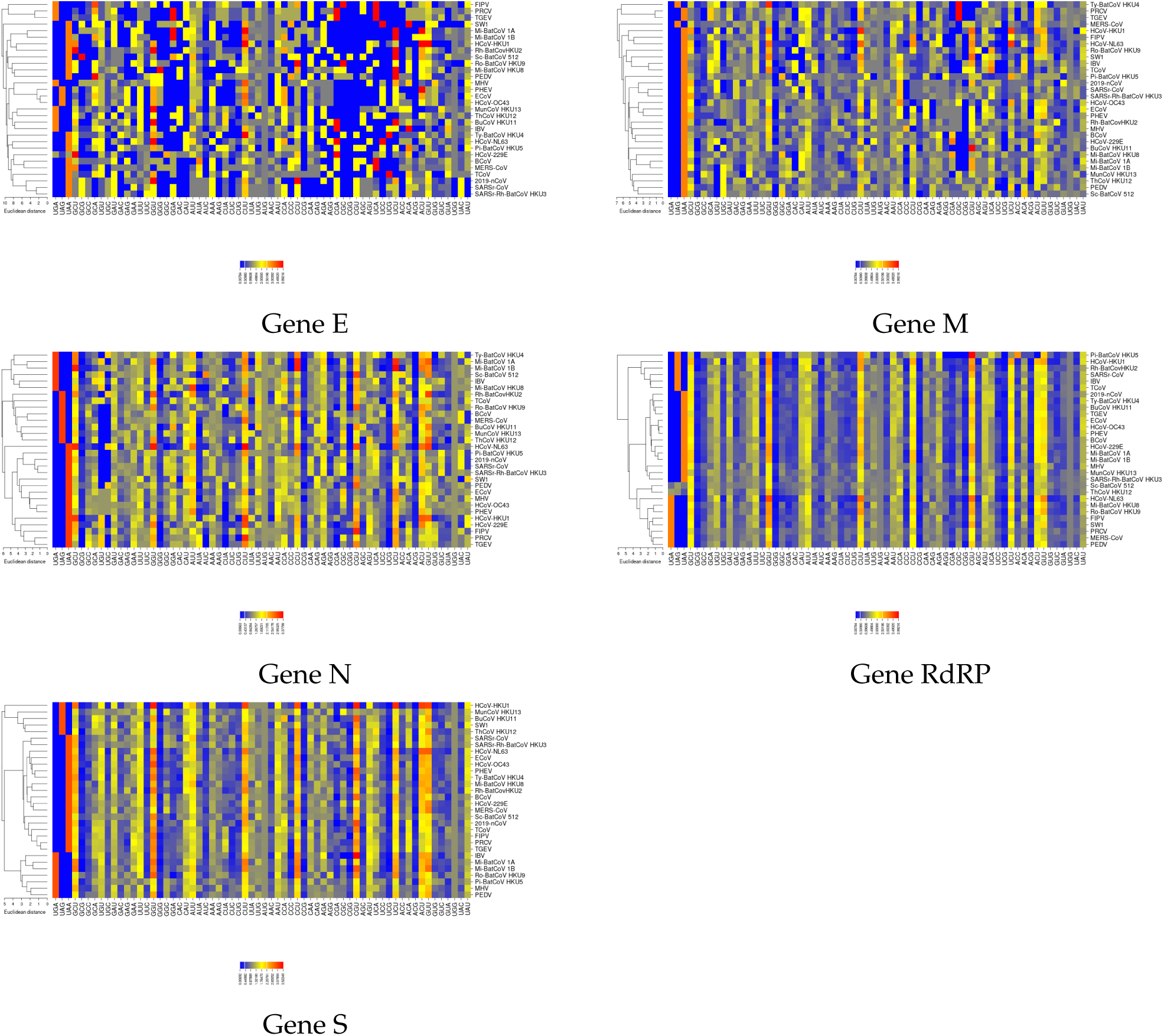
RSCU vectors of five different genes. Heatmaps confirm that the newly identified coronavirus 2019-nCoV Wuhan-Hu-1 sequence is closer to SARS and HCoV 229E, except for gene RdRP and gene S.

### 2.6. The ENC plot analysis of individual genes of 2019-nCoV

To further investigate which factors account for the low codon usage bias of the different genes in coronavirus, we analyzed the relationship between the ENC value and the percentage of G or C in the third positioncodons (GC3s). In Figure 4, we show separately the ENC-plots obtained for the five genes (M, N, S, E, and RdRP), together with the Wright’s theoretical curve, corresponding to the case when the codon usage is only determined by the GC3s [29]. We would like to point out that if mutational bias, as quantified by GC-content in the generally neutral 3rd codon position, is the main factor in determining the codon usage among these genes, the corresponding point in the ENC-plot should lie on or just below Wright’s curve. In Figure 4, all distributions lie below the theoretical curve, an indication that not only mutational bias but also natural selection plays a non-negligible role in the codon choice in all genes. This is exemplified by the violinplots in Figure 5 of the distances between the genes and Wright’s theoretical curve. Genes N, S, and RdRP are more scattered below the theoretical curve than those of genes M and E, implying that in the latter the codon usage patterns are pretty consistent with the effects of mutational bias. Interestingly, data points corresponding to the gene N, which is the major viral structural component needed to protect and encapsidate the viral RNA, is localized around GC3=0.5 (See Figure 4). This means that the displacement under the curve most likely reflects the selective pressure exerted on this gene. Conservely, all other genes show a displacement towards lower values of GC3-content, thus corroborating our previously mentioned observation that coronaviruses tend to use codons that end with A and U (see section 2.3).

**Figure 4.**
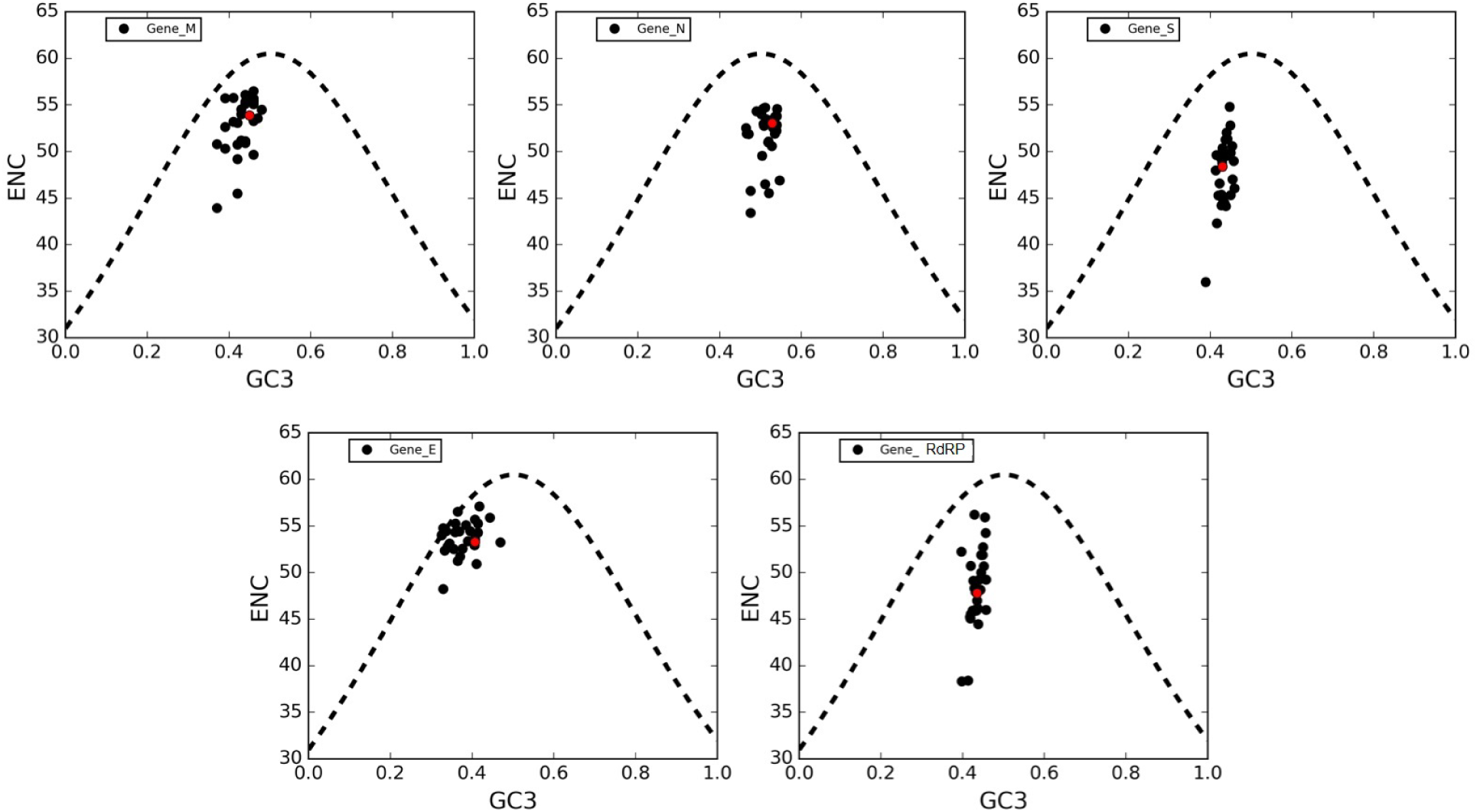
ENC-plots of genes M, N, S, E, RdRP. In these plots, each point corresponds to a single gene. The black-dotted lines in all panels are plots of Wright’s theoretical curve corresponding to the case of a CUB merely due to mutational bias (no selective pressure). Red dots represent 2019-nCoV genes.

**Figure 5.**
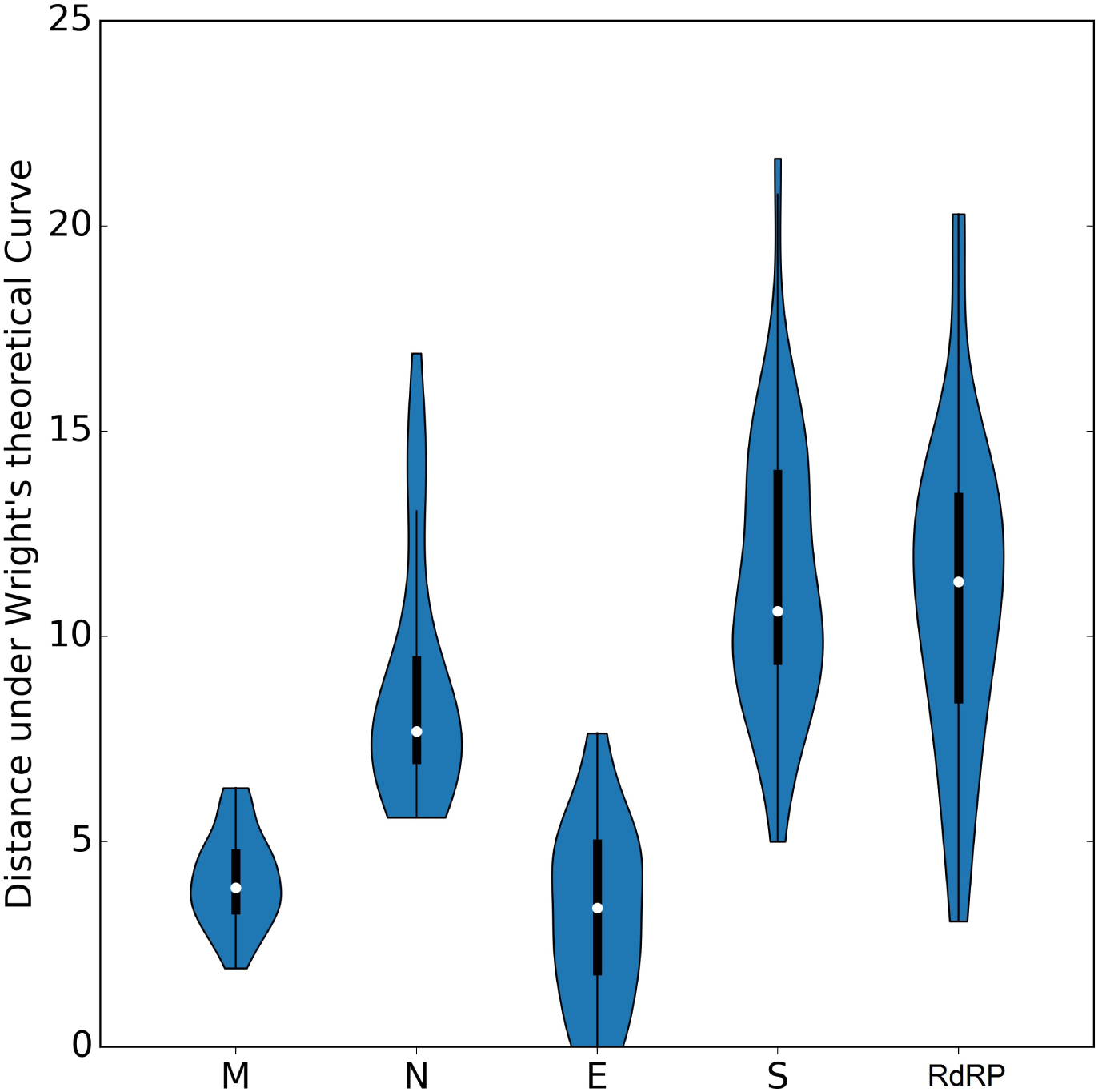
Distances from Wright’s curve. Box plots are calculated for each gene for values of distance from theoretical Wright’s curve. Genes N, S, and RdRP are more scattered below the theoretical curve than those of genes M and E, implying that in the latter the codon usage patterns are pretty consistent with the effects of mutational bias.

**Figure 6.**
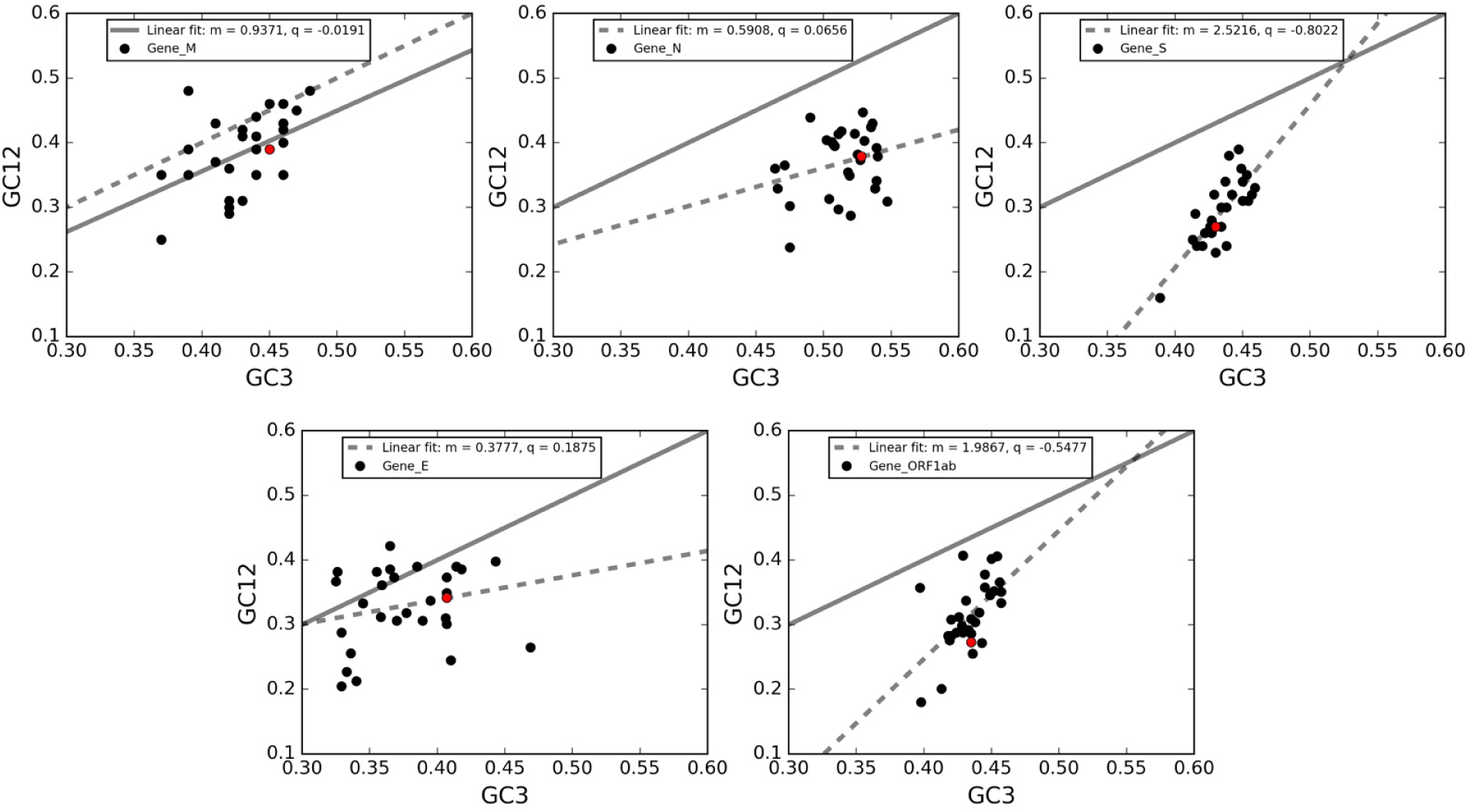
Neutrality plot of genes M, N, S, E, RdRP. In these plots, each point corresponds to a single gene in a virus. The solid black lines in all panels are the bisectors corresponding to the case of a CUB merely due to mutational bias (no selective pressure). The black-dotted lines are the linear regressions. Red dots represent 2019-nCoV genes.

### 2.7. Neutrality plot

A neutrality plot analysis was performed to estimate the role of mutational bias and natural selection in shaping the codon usage patterns of the five genes under investigation. In this plot, the average GC-content in the first and second positions of codons (GC12) is plotted against GC3s, which is considered as a pure mutational parameter. In Figure 5, we report the neutrality plots obtained for genes M, N, S, E, RdRP, together with the best-fit lines and the slopes associated with them.

The rationale to understand the results is that the wider is the deviation between the slope of the regression line and the bisector, the stronger is the action of selective pressure. All correlations are highly significant (Spearman correlation - *R*^2^ analysis, p-value<0.0001). By comparing the divergences between the regression lines and the bisectors in each panel, we reveal that the five genes here considered are subject to different balance between natural selection and mutational bias. Specifically, in line with the ENC-plot analyses, the genes S and RdRP present the largest deviations of their regression lines from the bisector lines and, therefore, a stronger action of natural selection. Conversely, the regression line for the gene M is closer to the bisector than the other genes, meaning that the this gene is the least one subject to the action of natural selection. Finally, the genes E and N are intermediate between the previous cases.

It is worth noting that almost all data points are located below the bisector lines, implying a selective tendency for a higher AT content in the first two codon positions than in the third one. In addition, both GC3 and GC12 are lower than 0.5, thus showing a general preference for A and T bases in all three codon positions. Interestingly, data points associated to gene M and E are closer to the bisector lines as compared to gene N, S, and RdRP. This observation means that the GC content in the first two codon positions tend to be in proportion to GC3 in gene M and E, and this partially explains the closeness of these two genes to the Wright theoretical curve in Figure 4.

### 2.8. Forsdyke plot

We analyze the rate of evolution that characterize these genes by confronting the nucleotide sequences of the newly emerging 2019-nCoV and their corrisponding protein sequences with those of other coronaviruses here considered. Each gene of 2019-nCoV was compared to the orthologous gene in one of the 30 coronaviruses to estimate evolutionary divergences. Each pair of homologous genes is represented by a point in the Forsdyke plot [8], which correlates protein divergence with DNA divergence. Each point in Forsdyke plots measures the divergence between pairs of orthologous genes in the two species, as projected along with the phenotypic (protein) and nucleotidic (DNA) axis. Thus, the slope is an estimation of the fraction of DNA mutations that result in amino acid substitutions in the speciation process between two species [8]. In Figure 7, we show separately for each gene the associated Forsdyke plot.

**Figure 7.**
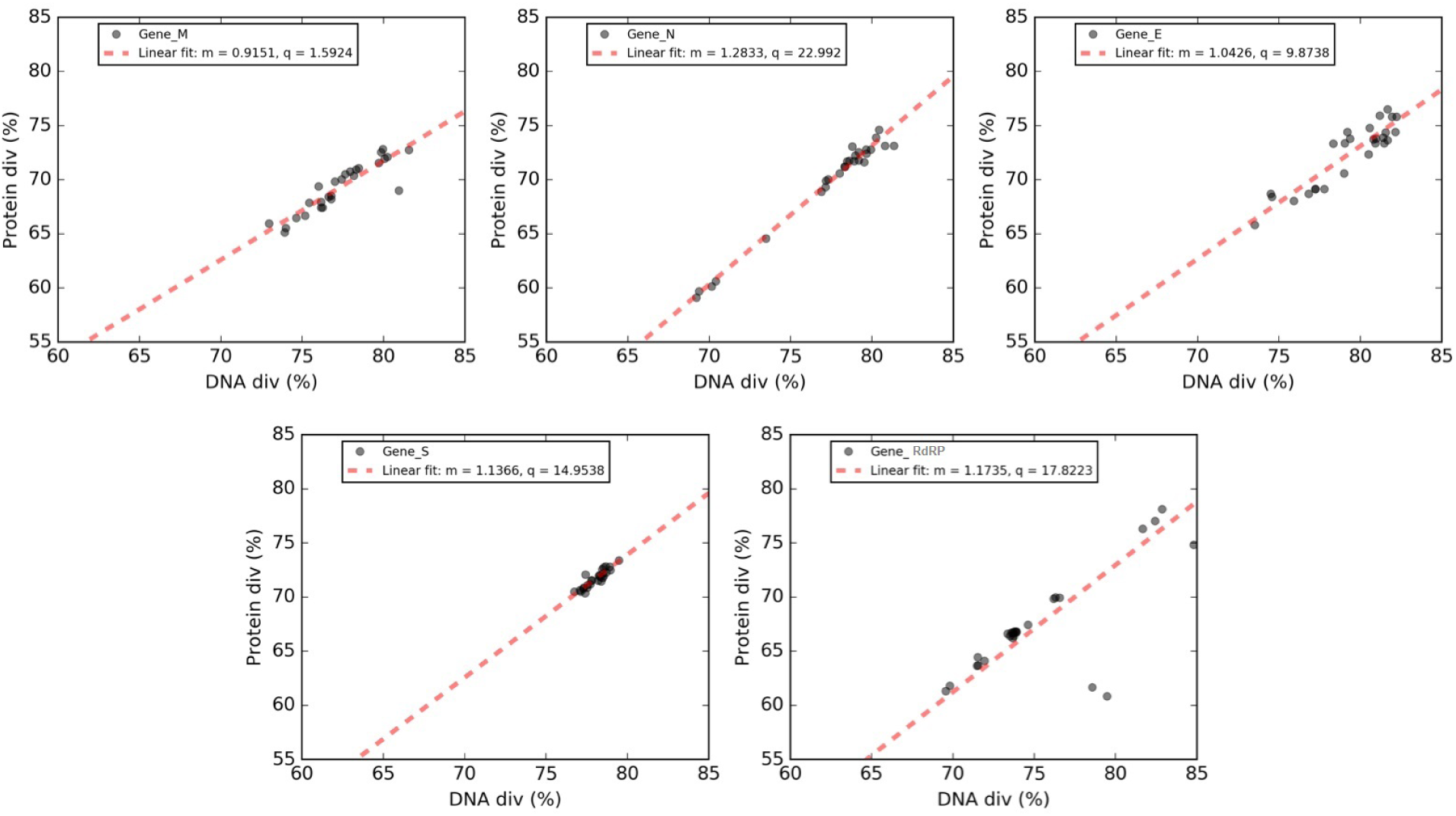
Forskdyke plots of genes M, N, S, E, RdRP. Phenotype (Protein div) vs. nucleotide (DNA div) sequence divergence between 2019-nCoV and homologous genes in the other coronaviruses. Each point corresponds to an individual gene. In each panel, we report the best-fit line in red, together with the associated values of the slope (m) and the intercept (q) in the legend.

Overall, protein and DNA sequence divergences are linearly correlated, and these correlations can be expressed through slopes and intercepts of the regression lines.

Genes M and E show quite low slopes, indicating that these proteins tend to evolve slowly by accumulating nucleotide mutations on their genes. Conversely, the steeper slopes for genes N, RdRP, and S indicate that these genes tend to evolve faster compared to other ones. A plausible explanation for this observation is that protein N due to its immunogenicity, has been frequently used to generate specific antibodies against various animal coronavirus, including in SARS [24]. The viral replicase polyprotein that is essential for the replication of viral RNA and, finally, the gene S encodes the protein that is responsible for the “spikes” present on the surface of coronaviruses. Taken together, our results suggest that the higher evolutionary rate observed for these proteins could represent a major obstacle to in the development of a antiviral therapeutics against 2019-nCoV.

## 3. Discussion

To investigate the factors leading to the 2019-nCoV and other viruses related to codon usage patterns, several analytical methods were used in our study. First, the RSCU value of the 2019-nCoV was calculated. Despite the high mutation rate that characterize 2019-nCoV, we do not show differences in codon usage between these genomes. Moreover, their associated vectors did not cluster for geographically positon, thus confirming the common origin of these genomes. The nucleotide composition confirm higher AT% content and low GC% content as it is common in RNA viruses such as Severe Acute Respiratory Syndrome (SARS) [**?**]. The results indicate also that codon usage bias exists and that the 2019-nCoV preferred codons almost all end in U. The codon usage bias was further confirmed by the mean ENC value of 51.9 (a value greater than 45 is considered as a slight codon usage bias due to mutation pressure or nucleotide compositional constraints). The same analysis is effectuated with CAI index, that it is calculated on the with RNA of codons expressed most often. This suggests that the RNA viruses with high ENC values (and low CAI) adapt to the host with various preferred codons. So a low biased codon usage pattern might allow the virus make use of several codons for each amino acid, and might be beneficial for viral replication and translation in the host cells. The ENC-plot analysis reveals that natural selection plays an important role in the codon choice of the five conserved viral genes here considered: RdRP, S, E, M, and N. However, genes N, S, and ORF1ab are more scattered below the theoretical curve than those of genes M and E, implying that in the latters the codon usage is under more strict control by mutational bias. According with this analysis, the genes S and RdRP present the largest deviations of their linear regressions from the bisectors lines in the neutrality plot analysis that implies a stronger action of natural selection. Conversely, the regression line for the gene M is closer to the bisector than the other genes, meaning that this gene is the least one subject to the action of natural selection. Finally, the genes E and N are intermediate between the previous cases.

To analyze the rate of evolution that characterize these genes by confronting the nucleotide sequences we use the Forsdyke plot thar correlates protein divergence with DNA divergence. The two proteins M and E show quite low slopes, indicating that these proteins tend to evolve slowly by accumulating nucleotide mutations on their genes. Conversely, the steeper slopes for gene N and RdRP and the gene S encoding for the S protein, which is responsible for the “spikes” present on the surface of coronaviruses, indicate that these three genes, and therefore their corresponding protein products, tend to evolve faster compared to the other two genes. Findings of the present study could be directed to be useful for developing diagnostic reagents and probes for detecting a range of viruses and isolates in one test and for vaccine development, using the information about codon usage patterns in these genes.

## 4. Materials and Methods

### 4.1. Sequence data analyzed

The complete coding genomic sequences of 306 isolates of nCov reported across the world to date, were obtained from GISAID (available at *https://www.gisaid.org/epiflu-applications/next-hcov-19-app/*) and NCBI viruses database accessed as on 17 March 2020.

Then the sequences are selected according to their geographical distribution, the isolation date, and the host species. The complete coding genomic sequences of 30 coronaviruses were downloaded from the National Center for Biotechnological Information (NCBI) (available at *https://www.ncbi.nlm.nih.gov/*). For each virus we consider in following alphabetical order genes: E, M, N, RdRP and S.

### 4.2. Nucleotide Composition Analysis

The diverse nucleotide compositional properties are calculated for the coding sequences of 30 CoV genomes. These compositional properties comprise the frequencies of occurrence of each nucleotide (A, T, G, and C), AU and GC contents, nucleotides G + C at the first (GC1), second (GC2), and third codon positions (GC3). To calculate these values we use an in-house Python script. We calculate, also, mean frequencies of nucleotides G + C at the first and the second positions (GC12).

### 4.3. RSCU

*RSCU* vectors for all the genomes were computed by using an in-house Python script, following the formula:

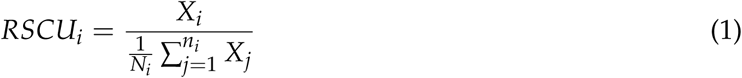

In the *RSCU*_*i*_ *X*_*i*_ is the number of occurrences, in a given genome, of codon i, and the sum in the denominator runs over its *n*_*i*_ synonymous codons. If the *RSCU* value for a codon *i* is equal to 1, this codon was chosen equally and randomly. Codons with *RSCU* values greater than 1 have positive codon usage bias, while those with value less than 1 have relatively negative codon usage bias [**?**]. *RSCU* heat maps were drawn with the CIMminer software [25], which uses Euclidean distances and the average linkage algorithm.

### 4.4. Effective Number of Codons Analysis

*ENC* is an estimate of the frequency of different codons used in a coding sequence. In general, *ENC* ranges from 20 (when each aminoacid is coded by just one and the same codon) to 61 (when all synonymous codons are used on an equal footing). Given a sequence of interest, the computation of *ENC* starts from *F*_*α*_, a quantity defined for each family *α* of synonymous codons (one for each amino acid):

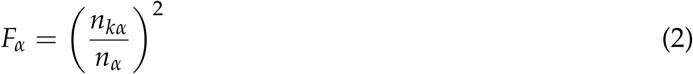

where *m*_*α*_ is the number of different codons in *α* (each one appearing 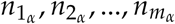 times in the sequence) and 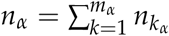.

*ENC* then weights these quantities on a sequence:

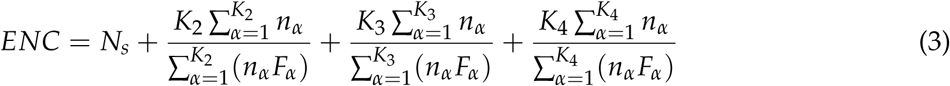

where *N*_*S*_ is the number of families with one codon only and *K*_*m*_ is the number of families with degeneracy *m* (the set of 6 synonymous codons for *Leu* can be split into one family with degeneracy 2, similar to that of *Phe*, and one family with degeneracy 4, similar to that, e.g., of *Pro*).

We have evaluated *ENC* by using the implementation in *DAMBE* 5.0 [31].

### 4.5. Codon Adaptation Index

Codon adaptation index *CAI* [**?**] is used to quantify the codon usage similarities between the virus and host coding sequences. The principle behind *CAI* is that codon usage in highly expressed genes can reveal the optimal (*i.e*., most efficient for translation) codons for each amino acid. Hence, *CAI* is calculated based on a reference set of highly expressed genes to assess, for each codon *i*, the relative synonymous codon usages (*RSCU*_*i*_) and the relative codon adaptiveness (*w*_*i*_):

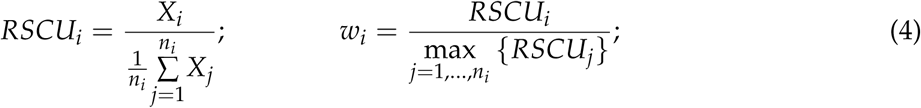

In the *RSCU*_*i*_, *X*_*i*_ is the number of occurrences of codon *i* in the genome, and the sum in the denominator runs over the *n*_*i*_ synonyms of *i*; *RSCU*s thus measure codon usage bias within a family of synonymous codons. *w*_*i*_ is then defined as the usage frequency of codon *i* compared to that of the optimal codon for the same amino acid encoded by *i*—*i.e*., the one which is mostly used in a reference set of highly expressed genes. The *CAI* for a given gene *g* is calculated as the geometric mean of the usage frequencies of codons in that gene, normalized to the maximum *CAI* value possible for a gene with the same amino acid composition:

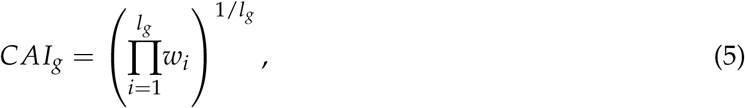

where the product runs over the *l*_*g*_ codons belonging to that gene (except the stop codon).

This index values range from 0 to 1, where the score 1 represents the tendency of a gene to always use the most frequently used synonymous codons in the host. The CAI analysis of these coding sequences are performed using *DAMBE* 5.0 [31]. The synonymous codon usage data of different hostes (human-Homo sapiens and other species) are retrieved from the codon usage database (available at: http://www.kazusa.or.jp/codon/).

To study the patterns of codon bias in the coronaviruses, we use Z-score values:

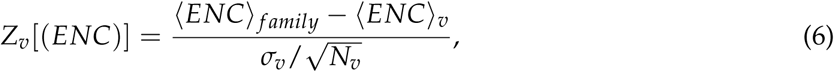

where ⟨*ENC*⟩ _*f amily*_ is the average of the ratio within a codon bias index in a family of a virus *v*, ⟨*ENC*⟩_*v*_ and *s*_*v*_ are the average value of *ENC* and its standard deviation over the whole virus *v*, and *N*_*v*_ is the number of virus in the family (we use the standard deviation of the mean as we are comparing average values).

The same Z-score is evalueted for codon bias index CAI.

### 4.6. ENC plot

ENC-plot analysis was performed to estimate the relative contributions of mutational bias and natural selection in shaping CUB of genes encoding for proteins that are crucial for 2019-nCoV: RdRP, the spike-surface glycoprotein (protein S), the small envelop protein (protein E), the matrix protein (M), and the nucleocapsid protein (N). The ENC-plot is a plot in which the ENC is the ordinate and the GC3 is the abscissa. Depending on the action of mutational bias and natural selection, different cases are discernable. If a gene is not subject to selection, a clear relationship is expected between ENC and GC3 [**?**]:

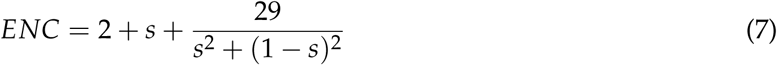

where *s* represents the value of *GC*_3_ [**?**]. Genes for which the codon choice is only constrained by mutational bias are expected to lie on or just below the Wright’s theoretical curve. Alternatively, if a particular gene is subject to selection, it will fall below Wright’s theoretical curve. In this case, the vertical distance between the point and the theoretical curve provides an estimation of the relative extent to which natural selection and mutational bias affect CUB.

To evaluate scatter of dots from theoretical Wright’s curve, we calculate the module of distance and we drawn box plot calculated with an in-house Python script.

### 4.7. Neutrality plot

We used a neutrality plot analysis [**?**] to estimate the relative contribution of natural selection and mutational bias in shaping CUB of five crucial genes in the research field aiming to develop a vacine against 2019-nCoV: M, N, S, RdRP, and E. In this analysis, the GC1 or GC2 values (ordinate) are plotted against the GC3 values (abscissa), and each gene is represented as a single point on this plane. In this case, the three stop codons (TAA, TAG, or TGA) and the three codons for isoleucine (ATT, ATC, and ATA) were excluded in calculation of GC3, and two single codons for methionine (ATG) and tryptophan (TGG) were excluded in all three (GC1, GC2, GC3) (Sueoka 1988). For each gene, we separately performed a Spearman correlation analysis between GC1 and GC2 with the GC3. If the correlation between GC1/2 and GC3 is statistically significant, the slope of the regression line provides a measure of the relative extent to which natural selection and mutational bias affect CUB of these genes (Sueoka 1999). In particular, if the mutational bias is the driving force that shapes CUB, then the corresponding data points should be distributed along the bisector (slope of unity). On the other hand, if natural selection also affects the codon choice of a family of genes, then the corresponding regression line should diverge from the bisector. Thus, the extent of the divergence between the regression line and the bisector indicates that the extent of codon choice due to the natural selection.

### 4.8. Forsdyke plot

To study the evolutionary rates of genes M, N, S, RdRP, and E, we performed an analysis by using our previously defined Forsdyke plot [8]. Each gene in 2019-nCoV (taken as a reference) was confronted with the homologous gene in one of the 30 coronaviruses considered in this analysis. Each pair of homologous genes is represented by a point in the Forsdyke plot, which correlates protein divergence with DNA divergence (see Methods in [8] for details). The protein sequences were aligned using Biopython. The DNA sequences were then aligned using the protein alignments as templates. Then, DNA divergences and protein divergences were assessed as axplained in Methods in [8] by counting the number of mismatches in each pair of aligned sequences. Thus, each point in Forsdyke plots measures the divergence between pairs of homologous genes in the two species, as projected along with the phenotypic (protein) and nucleotidic (DNA) axis.The first step in each comparison is to compute the regression line between protein vs. DNA sequence divergence in the Forsdyke plot getting values of intercept and slope for each variant of gene. To test whether the regression parameters associated with each variant are different or not, we have followed a protocol founded by F-statistic test, considering a p-value *≤* 0.05.

### 4.9. Phylogenetic analysis

To explore the evolutionary relationships among the four genera of coronaviruses, phylogenetic analysis of the full-length genomic sequences of the 30 CoVs listed in Table 1 was performed. The sequences were aligned with the usage of ClustalO [21], [22]. The resulting multiple sequence alignment was used to build a phylogenetic tree by employing a maximum-likelihood (ML) method implemented in the software package MEGA version 10. 1 [15]. ModelTest-NG [6] was used to select the best-fit evolutionary model of nucleotide substitution, that is, GTR + G + I. Bootstrap analysis (100 pseudo-replicates) was conducted in order to evaluate the statistical significance of the inferred trees.

## Author Contributions

For research articles with several authors, a short paragraph specifying their individual contributions must be provided. The following statements should be used “Conceptualization, all.; methodology, M.D, S.F.and A.P..; software, M.D and S.F.; validation, all.; formal analysis, M.D, S.F A.P.; investigation, M.D, S.F., A.P; data curation, M.D, S.F., A.P. ; writing–review and editing, all. All authors have read and agreed to the published version of the manuscript.’, please turn to the CRediT taxonomy for the term explanation. Authorship must be limited to those who have contributed substantially to the work reported.

## Conflicts of Interest

The authors declare no conflict of interest.

## Abbreviations

The following abbreviations are used in this manuscript:

MDPI: Multidisciplinary Digital Publishing Institute
DOAJ: Directory of open access journals
TLA: Three letter acronym
LD: linear dichroism

## Appendix A

**Appendix A.1.**
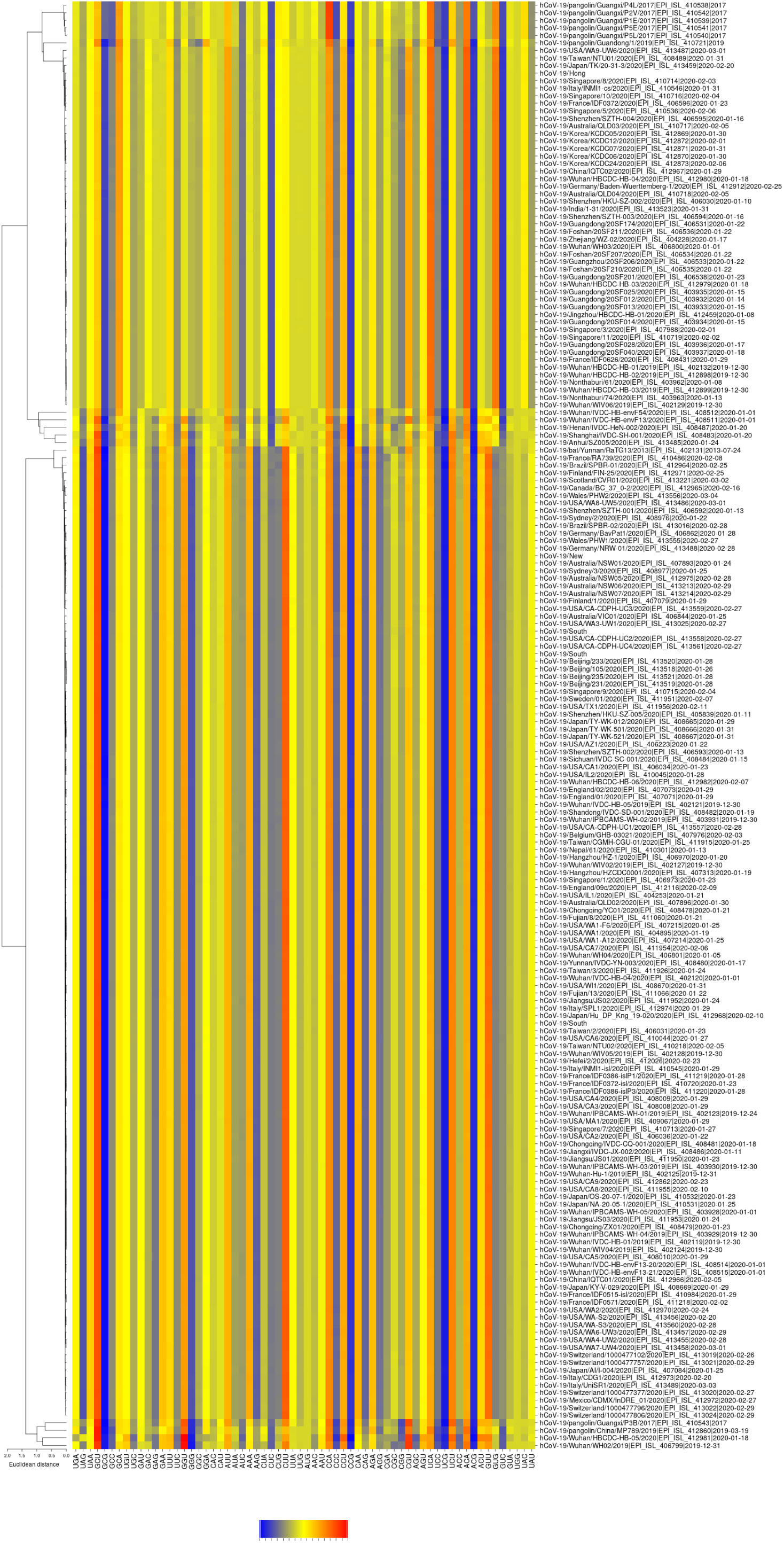

**Figure A2.**
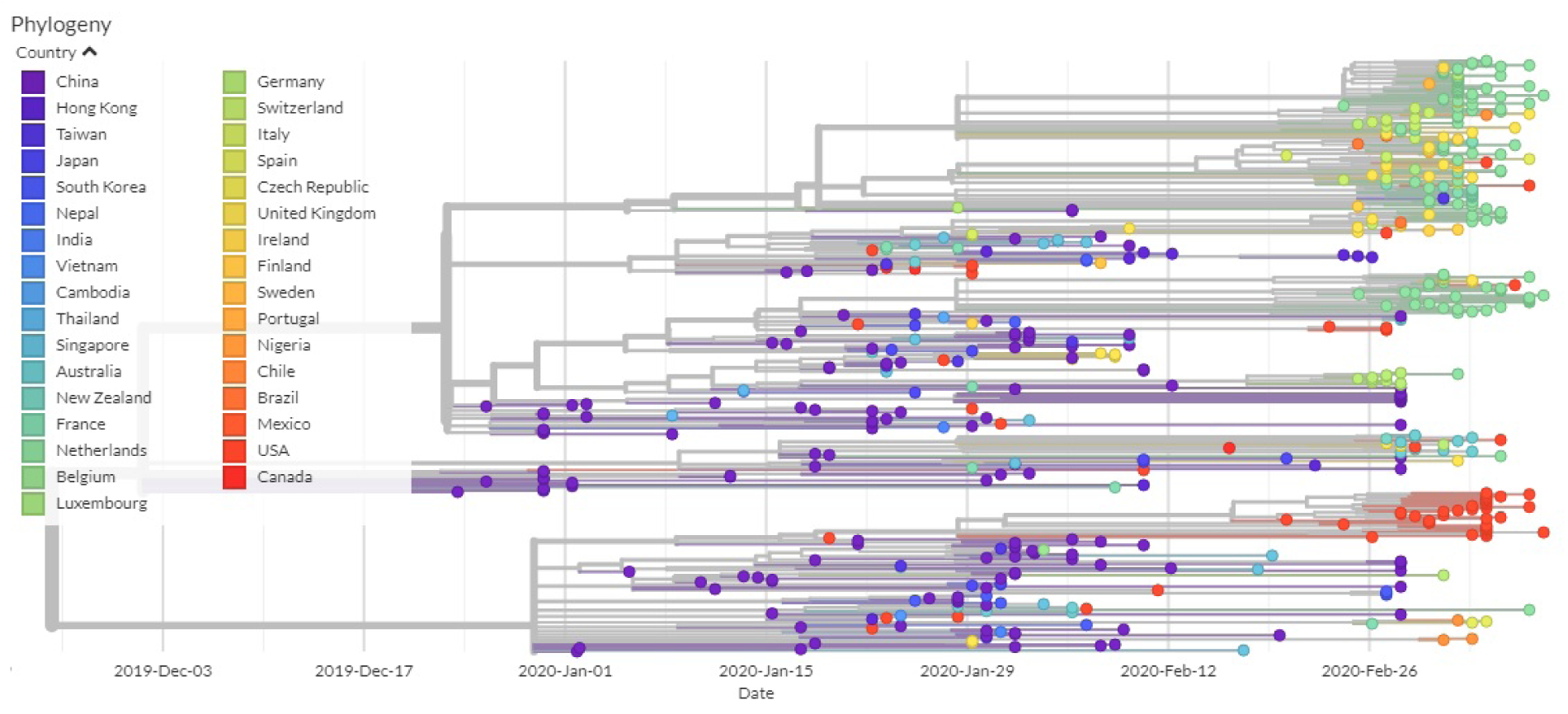
Phylogenetic tree from GIDAID.

**Figure A3.**
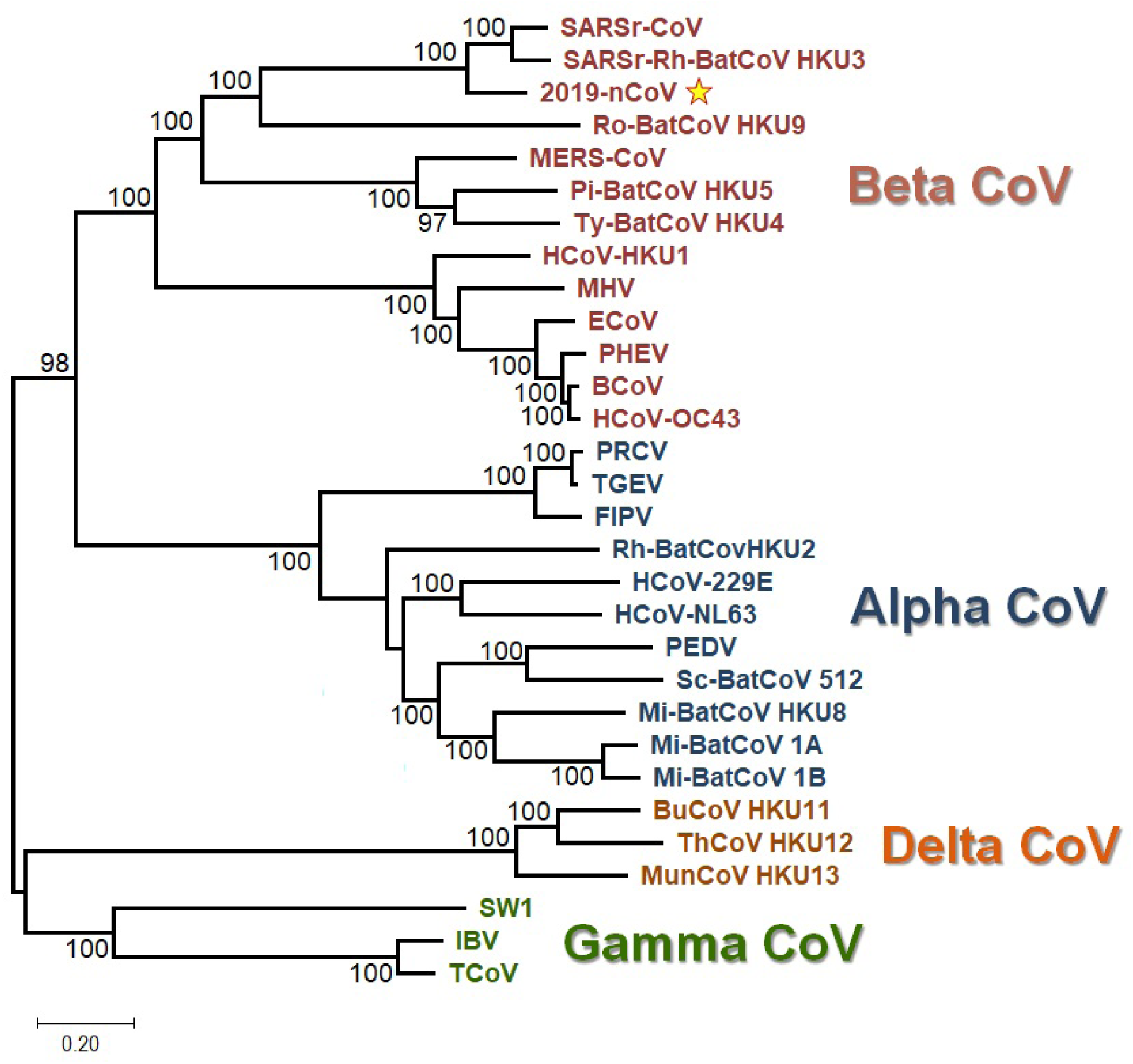
Unrooted ML-based tree of the 30 CoV genomic sequences. The four distinct color-coded clades correspond to the respective genera of CoVs. The 2019-nCoV sequence is indicated by a star. The branch lengths depict evolutionary distance. Bootstrap values higher than 50 are shown at the nodes. The scale bar at the lower left denotes the length of nucleotide substitutions per position.

## References

1. Belalov IS, Lukashev AN, Causes and Implications of Codon Usage Bias in RNA Viruses PLoS One. 2013; 8(2): e56642. Published online 2013, Feb 25. doi: 10.1371/journal.pone.0056642

2. Cavanagh D. The Coronavirus Surface Glycoprotein, the Coronaviridae Springer (1995), pp. 73–113

3. Ceraolo C., Giorgi FM. Genomic variance of the 2019-nCoV coronavirus, DOI: 10.1002/jmv.25700

4. Cao Y, Li L, Feng Z. et al. Comparative genetic analysis of the novel coronavirus (2019-nCoV/SARS-CoV-2) receptor ACE2 in different populations. Cell Discov 6, 11 (2020). https://doi.org/10.1038/s41421-020-0147-1

5. Chen Y, Xu Q, Yuan X, Li X. Analysis of the codon usage pattern in Middle East Respiratory Syndrome Coronavirus, Oncotarget, 2017, Vol. 8, (No. 66), pp: 110337–110349

6. Darriba D” Posada D, Kozlov AM, Stamatakis A” Morel B, Flouri T. ModelTest-NG: A New and Scalable Tool for the Selection of DNA and Protein Evolutionary Models., Mol Biol Evol. 2020 Jan 1;37(1):291–294. doi: 10.1093/molbev/msz189

7. Dilucca M, Cimini G, Giansanti A. Essentiality, conservation, evolutionary pressure and codon bias in bacterial genomes, Gene. 2018 Jul 15;663:178–188. doi: 10.1016/j.gene.2018.04.017. Epub 2018 Apr 18

8. Forcelloni, S, Giansanti A. Mutations in disordered proteins as early indicators of nucleic acid changes triggering speciation. Sci Rep 10, 4467 (2020). https://doi.org/10.1038/s41598-020-61466-5

9. Forni D, Cagliani R, Clerici M, Sironi M. Molecular Evolution of Human Coronavirus Genomes. Trends Microbiol. 2017; 25:35–48

10. Gorbalenya AE, Luis Enjuanes, John Ziebuhr, Eric J. Snijdera. Nidovirales: Evolving the largest RNA virus genome, Volume 117, Issue 1, April 2006, Pages 17–37 https://doi.org/10.1016/j.virusres.2006.01.017

11. Grantham R, Gautier C, Gouy M, Mercier R and Pavé A. Codon catalog usage and the genome hypothesis, Nucleic Acids Research, vol. 8, no. 1, pp. r49–r62, 1980

12. Gu W., Zhou T. Analysis of synonimous codon usage in SARS Coronavirus and other viruses in the Nidovirales, Virus Res. 2004 May;101(2):155–61

13. Jenkins GM and Holmes EC. The extent of codon usage bias in human RNA viruses and its evolutionary origin, Virus Research, vol. 92, no. 1, pp. 1–7, 2003

14. Ji W, Wang W, Zhao X, Zai J, Li X. Cross-species transmission of the newly identified coronavirus 2019-nCoV. J Med Virol. 2020;92: 433–440. https://doi.org/10.1002/jmv.25682

15. Kumar S Stecher G, Li M, Knyaz C, Tamura K. MEGA X: Molecular Evolutionary Genetics Analysis across Computing Platforms. Mol Biol Evol. 2018 Jun 1;35(6):1547–1549. doi: 10.1093/molbev/msy096

16. Lia G, Wang H, Wanga S, Xinga G, Zhanga C, Zhanga W. Insights into the genetic and host adaptability of emerging porcine circovirus, VIRULENCE 2018, VOL. 9, NO. 1, 1301–1313 https://doi.org/10.1080/21505594.2018.1492863

17. Neuman BW, Kiss G, Kunding AH, Bhella D, Baksh MF, Connelly S. A structural analysis of M protein in coronavirus assembly and morphology J. Struct. Biol., 174 (2011), pp. 11–22

18. Ruch T, Machamer C. The coronavirus E protein: assembly and beyond Viruses, 4 (2012), pp. 363–382

19. Sheihk. Analysis of preferred codon usage in the coronavirus N genes and their implications for genome evolution and vaccine design, Journal of Virological Methods 277 (2020) 113806

20. Siddell SG, Ziebuhr J, Snijder EJ. Coronaviruses, Toroviruses, and Arteriviruses. Topley and Wilson’s Microbiology and Microbial Infections (2005)

21. Sievers F, Higgins DG. Clustal omega. Curr Protoc Bioinformatics. 2014 Dec 12;48:3.13.1-16. doi: 10.1002/0471250953.bi0313s48

22. Sievers F, Higgins DG. Methods Mol Biol. 2014;1079:105–16. doi: 10.1007/978-1-62703-646-76 Clustal Omega, accurate alignment of very large numbers of sequences

23. Sueoka N. Directional mutation pressure and neutral molecular evolution. Proc Natl Acad Sci U S A. 1988; 85:2653–2657

24. Timani KA, Ye L, Ye L, Zhu Y, Wu Z, Gong Z. Cloning, sequencing, expression, and purification of SARS-associated coronavirus nucleocapsid protein for serodiagnosis of SARS, J Clin Virol. 2004 Aug;30(4):309–12.

25. Weinstein JN, Myers TG, O’Connor PM, Friend SH, Fornace AJ Jr, Kohn KW, Fojo T, Bates SE, Rubinstein LV, Anderson NL, Buolamwini JK, van Osdol WW, Monks AP, Scudiero DA, Sausville EA, Zaharevitz DW, Bunow B, Viswanadhan VN, Johnson GS, Wittes RE, Paull KD (1997) An information-intensive approach to the molecular pharmacology of cancer. Science 275(5298):343–9

26. Woo PC, Huang Y, Lau SK, Yuen KY. Coronavirus genomics and bioinformatics analysis. Viruses 2010 Aug;2(8):1804–20. doi: 10.3390/v2081803. Epub 2010 Aug 24. PMID: 21994708; PMCID: PMC3185738

27. Woo PC, Wong BH, Huang Y, Lau SK, Yuen KY. Cytosine deamination and selection of CpG suppressed clones are the two major independent biological forces that shape codon usage bias in coronaviruses, Virology 2007, 369, 431–442

28. Woo PC, Lau Y, Huang Y, Yuen KY. Coronavirus Diversity, Phylogeny and Interspecies Jumping. Experimental Biology and Medicine, 234(10), 1117–1127.(2009). https://doi.org/10.3181/0903-MR-94

29. Wright F. The ‘effective number of codons’ used in a gene, Gene. 1990 Mar 1;87(1):23–9

30. Wu A, Peng Y et al. Genome Composition and Divergence of the Novel Coronavirus (2019-nCoV) Originating in China Cell Host Microbe, Cell Host Microbe. 2020 Mar 11;27(3):325–328. doi: 10.1016/j.chom.2020.02.001. Epub 2020 Feb 7

31. Xia X, DAMBE5: a comprehensive software package for data analysis in molecular biology and evolution., Mol Biol Evol. 2013 Jul;30(7):1720–8. doi: 10.1093/molbev/mst064. Epub 2013 Apr 5

